# Data-Driven Mathematical Approach for Removing Rare Features in Zero-Inflated Datasets

**DOI:** 10.1101/2023.03.11.532198

**Authors:** Adrian N Ortiz-Velez, Scott T Kelley

**Affiliations:** Biological and Medical Informatics Program, San Diego State University, San Diego, CA, USA; Department of Biology, San Diego State University, San Diego, CA, USA

## Abstract

Sparse feature tables, in which many features are present in very few samples, are common in big biological data (e.g., metagenomics, transcriptomics). Ignoring the problem of zero-inflation can result in biased statistical estimates and decrease power in downstream analyses. Zeros are also a particular issue for compositional data analysis using log-ratios since the log of zero is undefined. Researchers typically deal with zero-inflated data by removing low frequency features, but the thresholds for removal differ markedly between studies with little or no justification. Here, we present CurvCut, a data-driven mathematical approach to zero-inflated feature removal based on curvature analysis of a “ball rolling down a hill”, where the hill is a histogram of feature distribution. These histograms typically contain a point of regime change, a discontinuity with a sharp change in the characteristics of the distribution, that can be used as a cutoff point for low frequency feature removal that considers the data-specific nature of the feature distribution. Our results show that CurvCut works well across a variety of biological data types, including ones with both right- and left-skewed feature distributions, and rapidly generates clear visual results allowing researchers to select data-appropriate cutoffs for feature removal.

## Introduction

Advancement in next-generation sequencing (NGS) technology has made it possible to detect thousands of species, genes, transcripts, or polymorphisms in samples, including ones at very low frequency [1]. This results in sparse feature tables dominated by zeros because many of the features are detected in only a few samples. These zero-inflated datasets have an excess of zeros in feature columns more than expected under a Poisson count distribution [2]. Ignoring the problem of zero-inflation can cause overdispersion and decrease power in downstream analyses [2, 3] or result in biased statistical estimates [4].

Zero-inflated datasets can be problematic for many types of statistical data analysis [4], including compositional data analysis (CoDA) methods [5, 6]. CoDA methods, originally developed for the geological sciences, have been increasingly applied to multiomics analysis (e.g., marker-gene libraries, transcriptomics, and metagenomics) due to their unavoidable compositionality [6–8]. CoDA approaches involve some sort of log-ratio transformation (e.g., centered log-ratio, isometric log-ratio), and therefore require replacement of all zeros in feature tables since the log of zero is undefined. Zero-replacement methods have been developed for this purpose [5, 6] but these methods do not work well with zero-inflated data. To overcome this limitation, researchers typically remove low-frequency features prior to zero-replacement. However, the process of feature removal is usually accomplished by setting an arbitrary threshold of percent presence among samples (i.e., only retain features present in at 10% of samples) and there appears to be no rule or consistent approach across studies for identifying that threshold. Thresholds for feature removal have been set at 10% [9], ≥1% in at least one sample [10], 85% [11, 12], or not reported or ignored [13–16]. While setting a percentage threshold for feature retention seems appealing, such a process does not consider the sparsity of individual datasets.

Here, we present a simple mathematical approach for selecting dataset-specific feature removal to reduce zero-inflation in a consistent manner that analyzes histogram distributions of dataset feature sparsity. These histograms typically contain a point of regime change, namely a discontinuity on the histogram with a sharp change in the characteristics of the distribution, and we detect this regime change using an implemented curvature analysis based on the idea of a ball rolling down a hill (Figure 1). We tested our approach using four different NGS feature tables generated from two small subunit ribosomal RNA (16S rRNA) amplicon datasets, one shotgun metagenomics dataset, and one single nucleotide polymorphism dataset. Our results show the curvature approach provides data-driven feature removal recommendations that consider the unique feature distributions of a given dataset.

**Fig 1.**
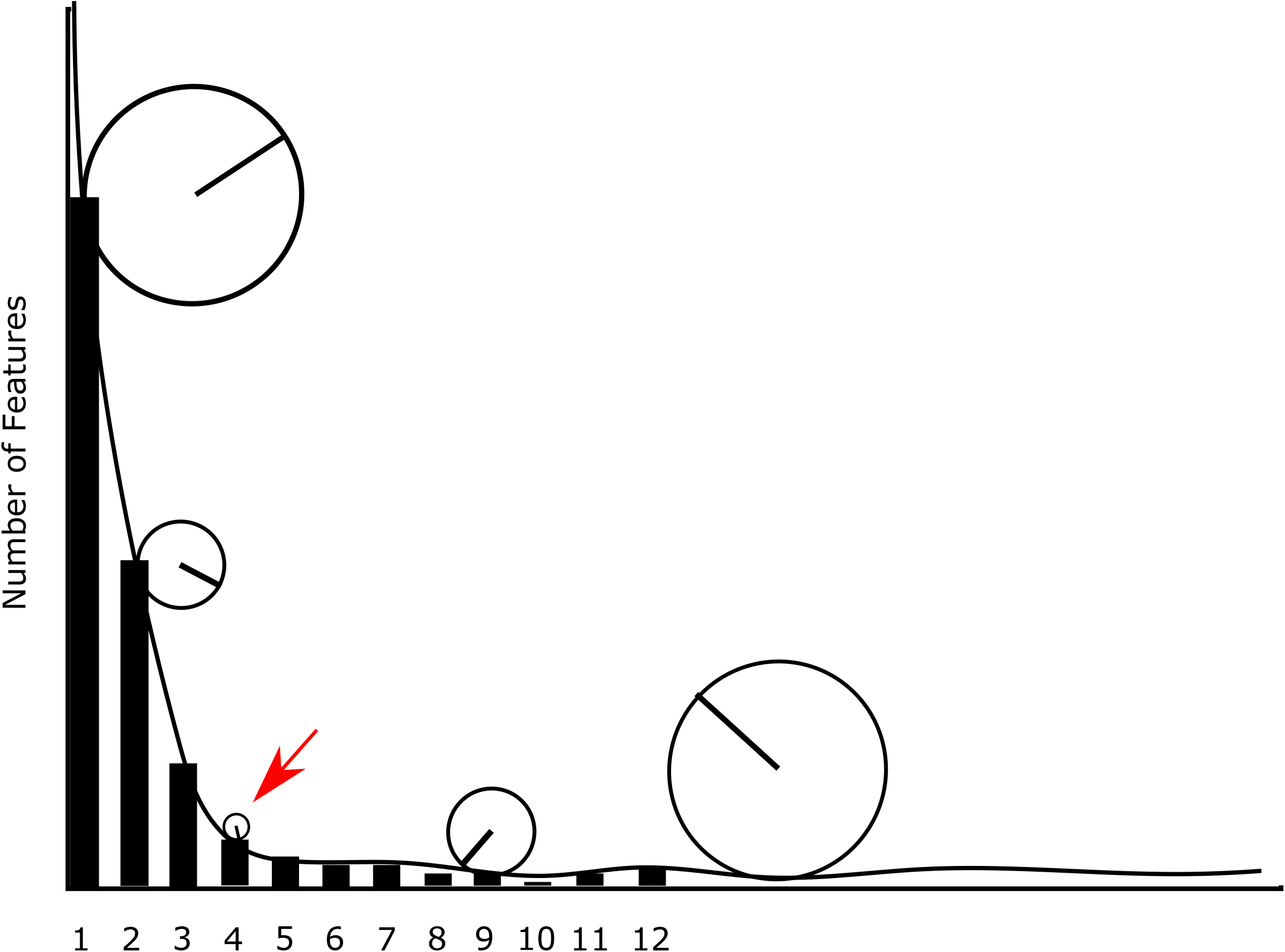
Artistic representation of the mathematical model. The histogram represents a hypothetical plot of features present per sample. The heights of the bars indicate the number of features (e.g., sequence variants, genes, single nucleotide polymorphisms) present in X samples. For example, the leftmost bar on the histogram represents features present in only one sample. To detect the regime change, we visualized the problem as a ball rolling down the hill of features. The radius of the ball decreases proportionally to the change in the height of the features until it reaches a minimum at the point of regime change when the path of the ball reaches the maximum curvature (i.e., when the ball is the smallest). Then after the regime change the ball increases again proportional to lack of change in the height of the feature as they reach steady values.

## Materials and methods

### Model Derivation

To create a mathematical model to detect the regime change in the zero-count distribution, we visualized the problem as a ball rolling down a hill of the feature distribution. In our model, the radius of the ball decreases proportionally to the height of the features (as it rolls down the hill) until it reaches a minimum at the point of ‘regime change’ when it shifts between characteristics of distributions (Figure 1). This allows detection of features that are exponentially decreasing. To generate the feature distributions, we counted the zeros in each column to make a histogram of bin size=1 for the zeros in the count table.

The first step in our approach is to create an accumulated zero count from a feature histogram. This is done because accumulation is characteristically monotonically increasing, so it is easier to look for a maximal change in the curvature across the log transformed zero count cumulative mass function. First, the zero count cumulative mass is calculated as

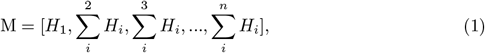

where H is the histogram array and n is the number of features. Next, the data are log transformed to maximize changes more than an order of magnitude and minimize changes less than an order of magnitude across the cumulative mass function. The log transform equation is

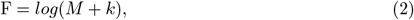

where F is the log transformed array of M and k is a constant (default k: 100) used to minimize small changes across M. To perform the final curvature analysis, we used CubicSplines from SciPy [17] to create a continuous piecewise polynomial function from the discontinuous histogram array

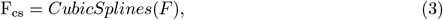

where F[cs] is the continuous piecewise function. To find the curvature across the cubic spline, we implemented the curvature equation

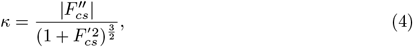

where K is the curvature and F”cs is the second derivative and F’ is the first derivative of the cubic spline Fcs. This has many local maxima so we identify the last maxima where

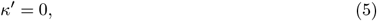

and we plot for the histogram C with the diagnosed cut-off for the user discernment [4].

### Implementation and Datasets

We implemented our curvature analysis method, called CurvCut, as a command line Python program (https://github.com/aortizsax/curvcut) and tested it on four datasets: two 16S datasets, one metagenomic count table dataset, and one HIV site frequency spectrum (SFS) dataset. The program was implemented in Python 3.7 using Python packages Pandas 1.3.3 [18], Numpy 1.19.2 [19], Matplotlib 3.3.4 [20], and SciPy 1.7.1 [17]. Scripts and the test datasets analyzed in this paper can be found at github.com/aortizsax/curvcut. The first two datasets, 16S and metagenomic count tables, come from a periodontal study [21]. The raw reads for the periodontal 16S rRNA sequences, and metagenomic OTUs classified by Kraken [22], were published previously [21, 23, 24]. The second 16S SV dataset comes from a built environment (BE) study [25].The HIV histogram was created by making a SFS graph of a multiple sequence alignment of Pol genetic sequence data with 100 sequences and 4339 bases long collected from NCBI BLASTN [26–35]. In this histogram, the heights of the bars represent the number of polymorphisms detected in one sequence(singletons), two sequences(doubletons), etc (polytons).

## Results and Discussion

Our results show that our data-driven modeling approach CurvCut identifies points of regime change across a variety of dataset types. CurvCut rapidly suggests a cutoff based on the distribution of features presence rather than on an arbitrary cutoff that does not consider the characteristics of the data (Figure 1). After curvature analysis, the recommended cutoff removes those features that could contribute to overdispersion in downstream analysis. The cutoff value is data-driven in the sense that the cutoff is entirely dependent on the feature distribution. The recommended cutoff value differs by dataset as expected by the clear differences in feature distributions (Figure 2). While most of the cutoffs suggested by our analysis were features that were in very few samples (5 or fewer), there was one dataset with a very high feature cutoff recommendation (features in 45 samples or less; Figure 2B). The importance of the cutoff being data-driven is very apparent when considering what would happen if the same cutoff was used for all the data sets in Figure 2. For example, a cutoff of features present in 3 or fewer samples would be appropriate for the allelic dataset (Figure 2D) but would leave many zeros in the 16S and metagenomics datasets (Figure 2A and B).

**Fig 2.**
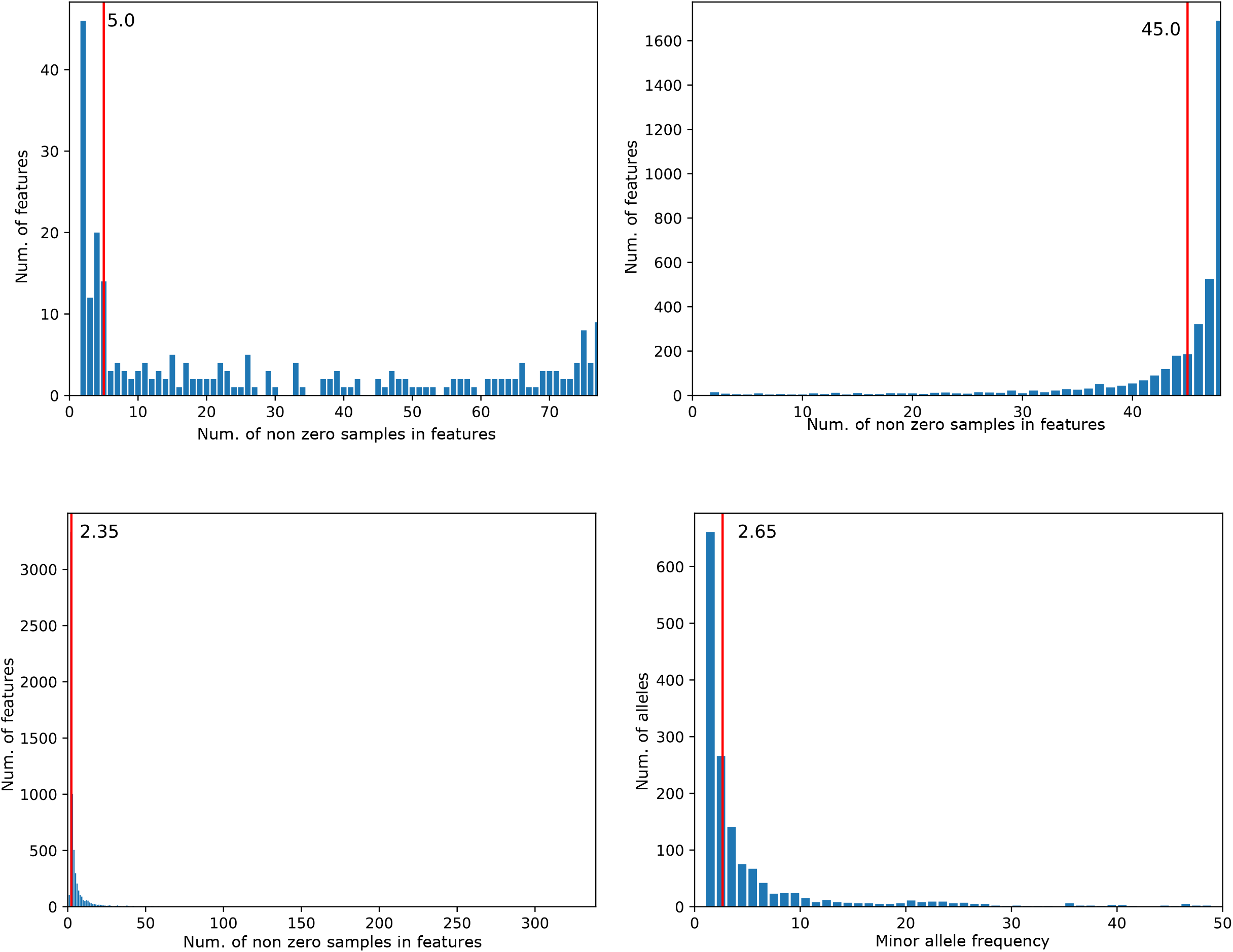
Curvature analysis of four datasets. The vertical red lines indicate the feature trimming cutoff based on our model. A) 16S periodontal data contained 76 samples of 247 OTU features, B) Metagenomic data contained 23 samples of 3,830 bacterial metagenomic features C) 16S built environment data containing 98 samples of 945 SV features, and D) HIV SFS data generated from an MSA of 100 Pol protein sequences, 4339 bases long.

A closer look at the trimmed sequence found many of them to be spurious. For example, the periodontal 16S dataset analyzed in Figure 2A trimmed two unidentified species of *Treponema*, most of which were missing in many or all of the samples, while an identified species, *Treponema socranskii*, was identified in all samples. While most of the distributions were right-skewed, our approach also worked on the left-skewed metagenomics dataset (Figure 2B), which shows that our method can detect these regime changes at either end of the distribution. A closer look at the metagenomic dataset analyzed in Figure 2B found that the k-mer based algorithm used to determine the number of species per sample, Kraken, identified certain species very readily in basically all the samples but also made many seemingly spurious identifications of closely related species or strains. For example, Kraken identified *Campylobacter gracilis* in every sample, with counts ranging between 900 and 330,000 (Avg.=64,000). However, Kraken also identified 15 other species of *Campylobacter*, most of which were missing in many or all the samples. This helps explain the leftward skew of this distribution and suggests that a cutoff of features present in 45 samples or fewer out of 48 samples would remove many spurious results. While most of the datasets were sequence count tables, our approach also worked on an SFS histogram from an HIV dataset (Figure 2D), which shows that our method can detect these regime changes in many types of datasets and is not exclusive to count tables. A closer look at the SFS dataset analyzed in Figure 2D, our analysis trimmed 661 and 266 of singletons and doubletons, which would remove possible noise and limit overdispersion in downstream analyses.

In general, we suspect that our method works best with right-or left-skewed features distributions, i.e., in datasets in which many features are present in only a few samples, or the opposite where many features are in many samples. This is usually the case with metagenomics, transcriptomics, and allelic datasets. However, some datasets might have a bimodal distribution of features that does not conform to this clear regime pattern. Therefore, CurvCut also allows the user to visualize the cutoff value on the feature distribution, so that they can make an informed choice on how to choose an appropriate cutoff. Our method can be easily integrated into common pipelines (e.g., QIIME2 [36] or mothur [37]) or run separately on datasets before further analysis.

## Data Availability

The CurvCut Programming code, installation instructions, and datasets used in this paper, including the HIV Pol MSA and SFS data are available at https://github.com/aortizsax/curvcut.

## Acknowledgments

We thank members of the Kelley Lab for testing CurvCut on numerous datasets. We also thank A. Sethuraman for preparing the SFS data and V. Thackray for comments on the manuscript.

